# Change of information content due to natural selection in populations with and without recombination

**DOI:** 10.1101/2020.01.23.917724

**Authors:** Wolfgang A. Tiefenbrunner

## Abstract

Though evolution undoubtedly operates in accordance with the second law of thermodynamics, the law of disorder, during billions of years organisms of incredible complexity came into being. Natural selection was described by Darwin^2^ as a process of optimization of the adaptation to environment, but optimization doesn’t necessarily lead to higher intricacy. Methods of thermodynamics and thus of information theory could be suited for the examination of the increase of order and information due to evolution.

Here I explain how to quantify the increase of information due to natural selection on the genotype and gene level using the observable change of allele frequencies. In populations with recombination (no linkage), the change of information content can be computed by summing up the contributions of all gene loci and thus gene loci can be treated as independent no matter what the fitness-landscape looks like. Pressure of deleterious mutations decreases information in a linear way, proportional to fitness loss and mutation rate.

The information theoretical view on evolution might open new fields of research.

## Introduction

Life on earth exists at least since 3.5 billion years^10,11,22^ and since then evolution led successively to the creation of organisms of ever increasing complexity. Evolution, the interplay of random mutations and purposeful selection, turned out to be a process of enormous creativity, astonishing for an optimizing procedure—with the simple aim to increase adaption to the environment—as which natural selection, the “driving force” of evolution, was described by Darwin 1859^2^:

“Hence, as more individuals are produced than can possibly survive, there must in every case be a struggle for existence, either one individual with another of the same species, or with the individuals of distinct species, or with the physical conditions of life.”

Fisher 1930^6^, in his mathematical formulation of the “genetical theory of natural selection”, took over the description as a process of optimization as did Rechenberg 1973^21^ in his artificial evolution experiments. However, optimization of adaption doesn’t necessarily lead to increasing complexity or creativity and so it seems that an important part of what makes natural selection so special is currently not understood enough.

Since Boltzmann^1^ found that the second law of thermodynamics is a law of disorder it was obvious that the existence of life, or more generally of self organization, doesn’t fit well to a world where the law of entropy rules. Major steps on the way to the understanding of this seemingly contradictory fact were Onsager’s^18^ (and others) non equilibrium thermodynamics, the theorem of minimum entropy production by Prigogine^20^ and the discovery of dissipative structures, the formulation of synergetics by Haken^8^ and the theory of molecular selection by Eigen^5^. Obviously there must be a bridge across the gap between thermodynamics and selection theory. Or, even more engaged, it must be possible to show that the theory of natural selection is simply a part of thermodynamics.

One step in this direction was done by Tiefenbrunner 1995^26^ by treating selection as a sorting process. If evolution may enhance complexity, the purposeful part of it, selection, may increase information, which means decrease entropy. This has to be quantified. In this paper I came to the conclusion that the change of information per generation and individual due to natural selection is given by

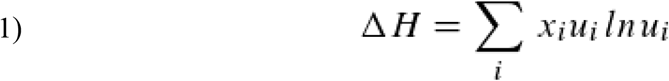

with

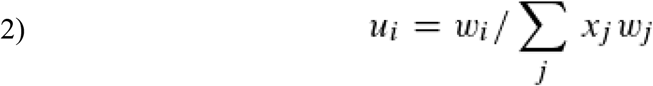

where ΔH is the change of information, w_i_ is the fitness of the i-th genotype in the population and x_i_ its frequency (actually it is the phenotype that survives and reproduces, or doesn’t, but it may be shaped by other forces than hereditary ones too. For the sake of simplicity we ignore the additional influence). Equation (1) is a good tool for the realm of theory but is not very useful in practice because the fitness is not immediately observable (but see ref. 7) or cannot be calculated from empirical data if the fitness-landscape is complex (Tiefenbrunner 1995 used equation (1) in a trial to generalize Eigen’s^5^ concept of an error threshold, a mere theoretical problem). However, in 1995 real time observations of evolutionary processes seemed to be not even a realistic aim for the nearer future. Since then things have changed insofar as now, thanks to new methods like whole-genome sequencing^7,12,25^ and bar coding^14,17^, it is relatively easy to measure gene and even genotype frequencies. To know how natural selection changes information may now be of empirical usefulness. Thus a new formulation of the connection between information and selection is necessary.

### Information change and natural selection

Suppose we have a population of N individuals that belong to i different genotypes (or phenotypes, the entity, selection deals with) and the frequency of the i-th one is ni. Then it follows that

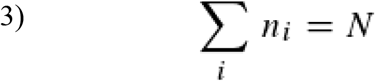

if x_i_ is the relative frequency of the i-th genotype, then:

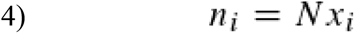

and

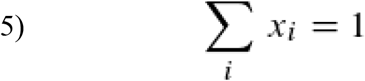

Let N’ be the number of specimens in the next generation, n_i_’ the frequency and x_i_’ the relative frequency of the i-th genotype in the next generation:

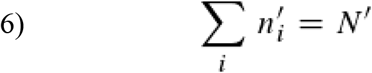

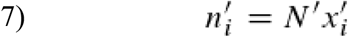

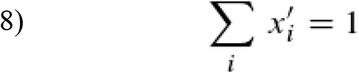

We separate selection and reproduction from each other in time and assume that selection occurs first. We suppose that after selection N’≪N. Of course after reproduction with propagation we have N’=N (constant size of population) and the next selection/reproduction-cycle may begin. Only selection but not reproduction shall change the frequencies of the genotypes. Now we may ask for the probability that the frequencies of the genotypes change within a generation in the observed way (that actually is a consequence of selection) by pure chance. According to Boltzmann^1^ (in the formulation of Planck^19^) this probability P of the observed composition (the one after selection) and entropy S are connected:

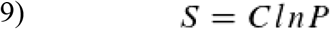

Because evolution is a successive process, discrete as we treat it, we will use ΔS, change of entropy per generation, instead of S. We simplify equation (9) by choosing C=1.

If we take individuals out of the population randomly (e.g. without selection), we get sequences of genotypes, where the frequency of the i-th genotype is n_i_’ (for all i), with the probability

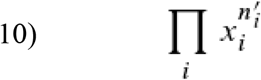

if n_i_ is large for all i, so that the probability of randomly picking out an individual of genotype i doesn’t change during the process. Of course we are not interested in sequences because it doesn’t matter when we pick out a specimen of genotype i but only how many we get finally. So we do not distinguish sequences as long as they do not differ in genotype frequencies. The number of sequences of the same genotype composition is given by

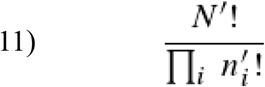

So the probability P to pick out randomly that composition which natural selection has picked out is:

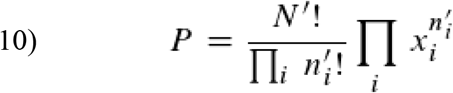

To simplify equation (10) we use Stirling’s approximation:

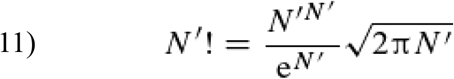

If we furthermore utilize the logarithm, small inaccuracy gets irrelevant, so that we can approximate:

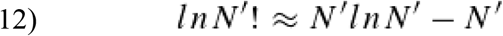

So we get

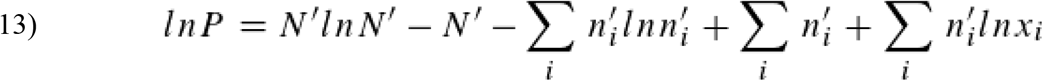

Because of equation (6) this is:

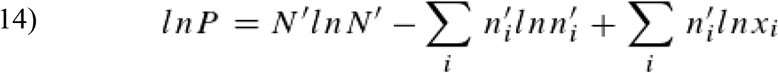

Using equation (7) we get:

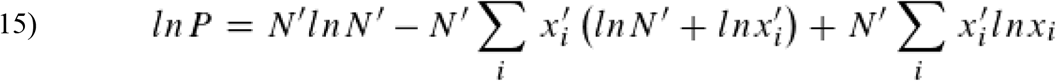

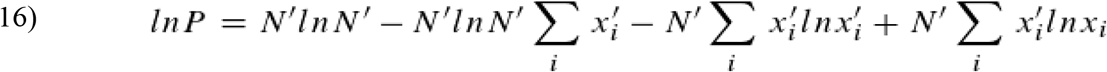

Equation (16) simplifies because of equation (8):

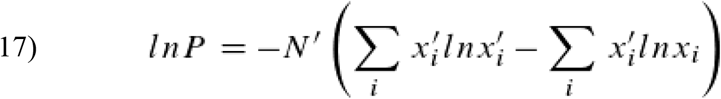

Because of equations (9) and (10) we see that entropy S of a composition is calculated as the difference between the entropy of an observed sequence and an expected one, expected under the assumption of randomness.

If we define ΔH as the information change per specimen and generation:

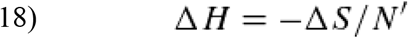

Finally we get:

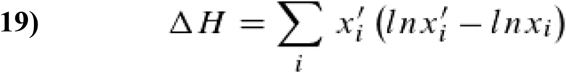

If x_i_’=0, then x_i_’(ln x_i_’–ln x_i_)=0. Equation (19) describes the information change due to natural selection as equation (1) does. But contrary to equation (1), equation (19) uses only observable parameters, the frequencies of genotypes in two consecutive generations and hence computation of ΔH is relatively easy.

Some of the assumptions we met to run the derivation of equation (19) were not very realistic, which makes equation (19) an approximation. More to this point later on.

What is the connection between equation (1) and equation (19)? For the derivation of equation (19) we didn’t need to know anything about the way natural selection works, contrary to the one of equation (1). Tiefenbrunner 1995^26^ interpreted selection as a sorting process (as we do here) and introduced a speciality of Thermodynamics, Maxwell’s demon, a being who sorts molecules by their kinetic energy (“a being whose faculties are so sharpened that he can follow every molecule in his course, and would be able to do what is at present impossible to us … Let us suppose that a vessel is divided into two portions A and B by a division in which there is a small hole, and that a being who can see the individual molecules opens and closes this hole, so as to allow only the swifter molecules to pass from A to B, and only the slower ones to pass from B to A. He will, thus, without expenditure of work raise the temperature of B and lower that of A, in contradiction to the second law of thermodynamics.”^15^). Maybe it’s more familiar to think of a speed trap that sorts cars by their velocity. This trap is special, like natural selection, because it’s fuzzy: higher speed means higher probability to be shot, but there is no level of velocity above which a photo is taken for sure. This fits to reality, for a less well adapted individual nevertheless may reproduce by chance as well as a very fit one may accidentally die without producing offspring. If we have i discrete speed classes, the probability of a car of class i to be shot is wi; and the proportional frequency of i-cars, from all the N cars that pass by, is x_i_. So we expect to find Nxiwi photos of i-cars in the resulting film (or directory that contains the digital photos) that consists altogether of NΣx_i_w_i_ =N’ photos. Hence the relative frequency x_i_’ of i-car photos in the film (or directory) is

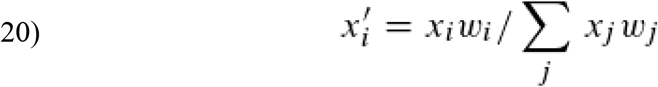

Equation (20) describes how the speed trap selection as a model for natural selection works. w_i_ then is the “fitness” of the car or genotype, respectively. From the equations (20) and (2) results:

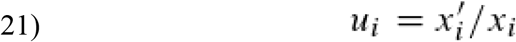

We may write equation (19) as

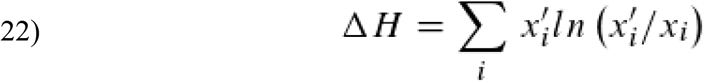

and finally within one step reach equation (1). So we see that equation (19) is more general than (1) because surely it would have been possible to use another model for selection. However, if we use equation (20) to describe selection, equation (1) results from (19).

Maybe it’s worth mentioning that selection makes reproduction necessary, an energetically costly procedure. The loss caused by selection (from N to N’) must be compensated from one generation to the next by reproduction with propagation (from N’ again back to N). To remain within our model, the loss of individuals that must be compensated depends on the mean fitness: the stronger the force of selection acts the energetically costlier is the compensatory propagation.

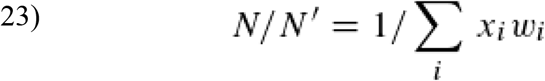

Equation (23) connects selection theory with the real, physical world, like the first law does with thermodynamics: selection costs energy (this is of course an idealized argumentation; individuals die anyway, even without selection).

### Information change, natural selection and recombination

Genetic recombination or reshuffling, the exchange of genetic material, is the great invention of life that is the centerpiece of sexual reproduction. There is occasionally also genetic recombination in asexual reproduction, but we do not deal with that here. If recombination occurs and there is no linkage between two gene loci, the frequency of the combinations of the alleles of the two loci (which is the haploid genotype in a two loci model) is given by the product of the frequencies of the alleles of the combination (the frequency of allele-combinations of any locus in diploid genotypes is given by the generalized Hardy-Weinberg law^9,27^).

In a population with recombination the information change due to natural selection behaves in a remarkable way. We find that

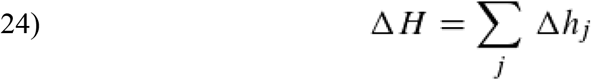

with

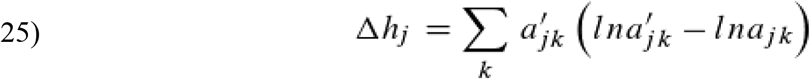

where a_jk_ is the relative frequency of the k-th allel of the j-th gene locus and a_jk_’ the proportional frequency of the same allele of the same locus in the next generation. Δh_j_ is the change of information content per generation due to selection of the j-th gene locus. If a_i_’=0, then a_i_’(ln a_i_’ – ln a_i_)=0.

One reason why equation (25) (together with (24)) is important is that the assumption for the derivation of equation (19) that all genotypes are frequent, isn’t a realistic one. On the gene level it is more likely that all alleles are frequent, which makes equation (25) the better approximation.

Of course,

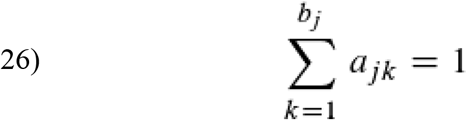

and

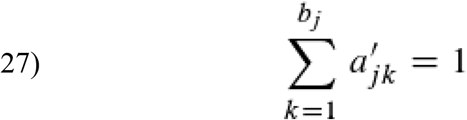

if there are b_j_ alleles in the j-th locus. Equation (24) remains valid, no matter what the fitness-landscape looks like. It is important to state this, because in every generation there is a haploid and a diploid phase with an assigned phenotype that is a target of natural selection so that the fitness-landscapes may be very manifold.

Now we have to prove equation (24). We start with a two loci (A_1^*^_, A_2^*^_), two alleles per loci (A_11_, A_12_, A_21_, A_22_) model. We have four haploid genotypes, X_1_, …, X_4_, that are represented by the gene combinations X_1_: A_11_A_21_, …, X_4_: A_12_A_22_. The past recombination frequency of the genotypes is

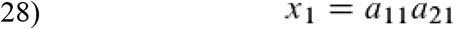

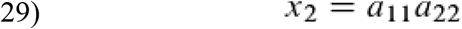

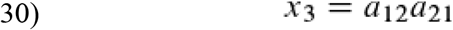

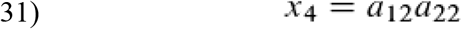

We use these to replace the x_i_ and x_i_’ in equation (19):

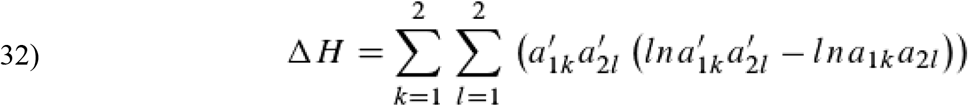

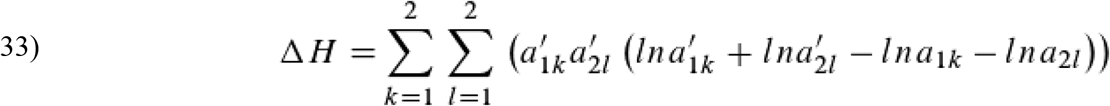

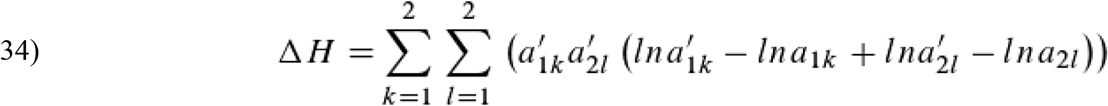

Which can be written as:

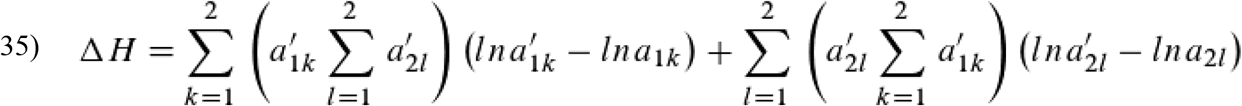

Because of equation (27) this can be simplified:

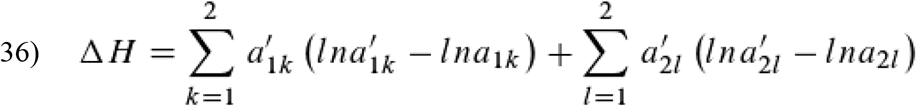

Finally we get

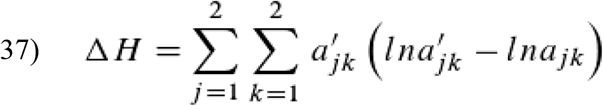

We can generalize this for more than two alleles per locus because the derivation remains the same

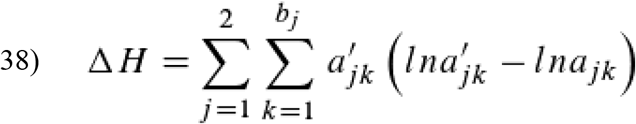

The generalization of the number of loci is more tricky. Thanks to recombination and what we already know we can treat two independent loci in mind as one with more pseudo-alleles—the combinations being the new alleles. To this pseudo-locus we add a new one and for these two equation 38 is valid. Now we repeat this approach again as often as we want, so that we finally come to the conclusion:

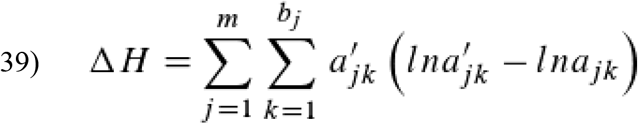

which is of course equal to equations (24) and (25).

Now let us look at some examples, especially those that point to the difference between populations with and without recombination. We use the two loci, two alleles per loci model. For selection we use equation (20), for recombination (28) to (31). Information change due to selection is computed with the aid of equation (19) on genotype level and on gene level using (37).

For the first example we assume additive fitness, e.g. the fitness of the genotype X_i_ is the sum of the fitness of the genes A_1k_ and A_2l_. (This is naturally an unrealistic assumption. Genes tend to adapt to their genetic environment as the existence of hybridisation zones and related phenomena show—see for instance the frequency dependent result of selection in Fig. 4 [top left] as possible basic for the explanation of hybridisation zones. Of course, thought experiments are in most cases simplifications).

**Fig. 1:**
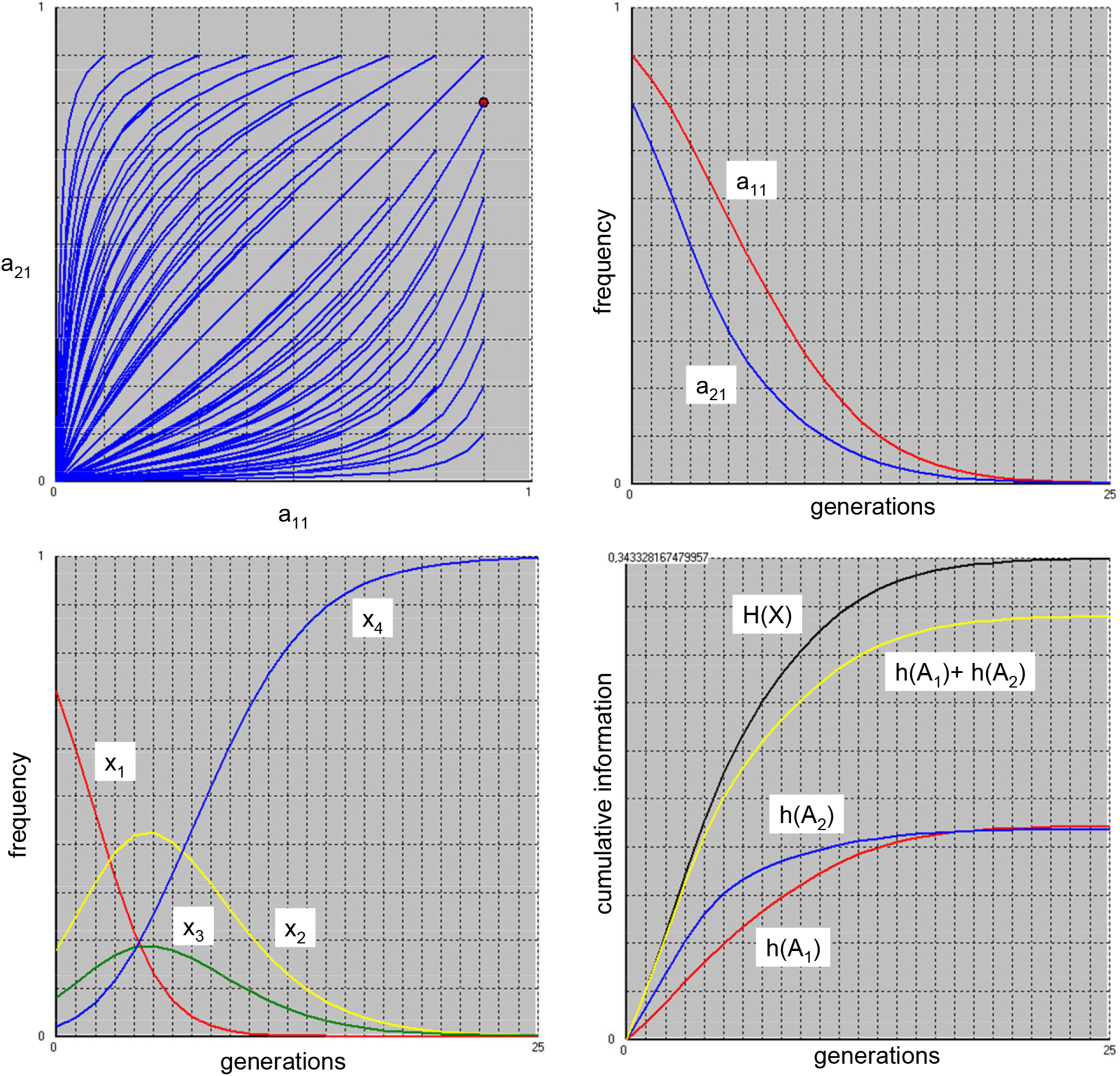
Selection **without recombination** in a two gene loci, two alleles per locus model **with additive fitness**. From left to right and top to bottom: [top left] change of frequency of A_11_ and A_21_ alleles shown against each other from different initial frequencies or [top right] shown in time (generations) from initial a_11_=0.9 and a_21_=0.8. Frequencies of the genotypes changing in time (generations) [bottom left]. Increase of information content due to selection on gene and genotype level[bottom right].

**Fig. 2:**
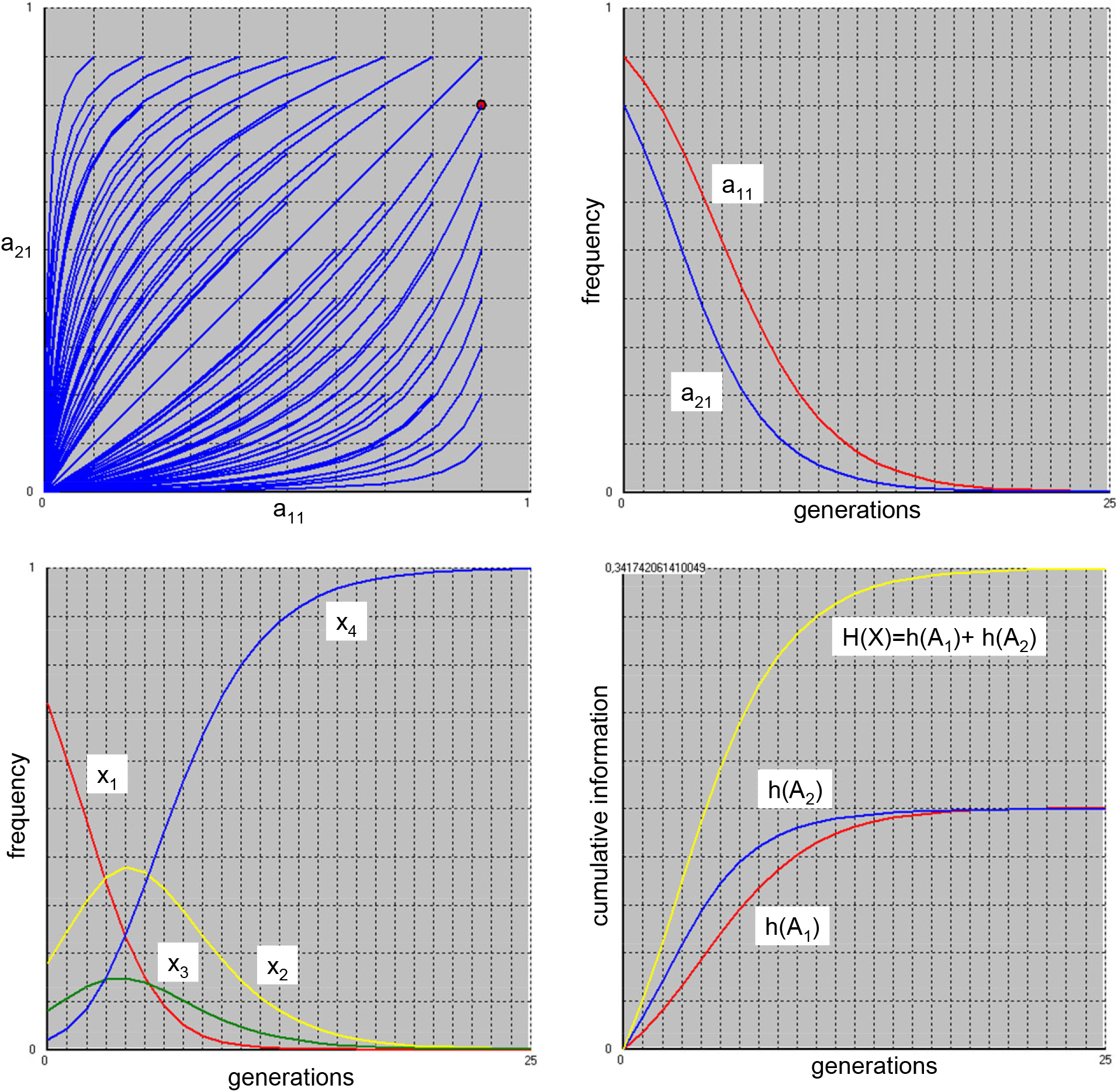
Selection **with recombination** in a two gene loci, two alleles per locus model **with additive fitness**. From left to right and top to bottom: [top left] change of frequency of A_11_ and A_21_ alleles shown against each other from different initial frequencies or [top right] shown in time (generations) from initial a_11_=0.9 and a_21_=0.8. Frequencies of the genotypes changing in time (generations) [bottom left]. Increase of information content due to selection on gene and genotype level [bottom right].

**Fig. 3:**
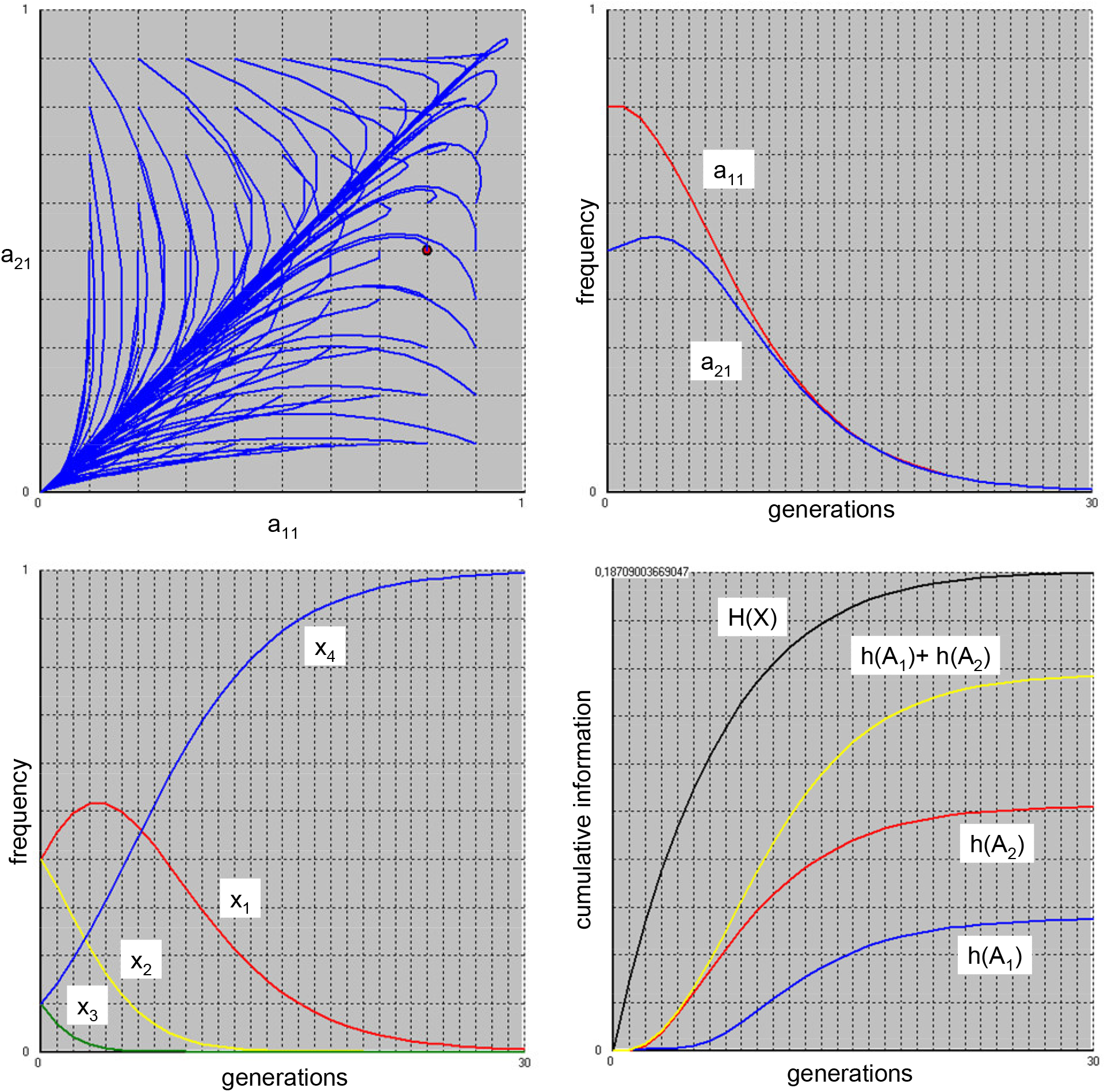
Selection **without recombination** in a two gene loci, two alleles per locus model **without additive fitness**. From left to right and top to bottom: [top left] change of frequency of A_11_ and A_21_ alleles shown against each other from different initial frequencies or [top right] shown in time (generations) from initial a_11_=0.8 and a_21_=0.5. Frequencies of the genotypes changing in time (generations) [bottom left]. Increase of information content due to selection on gene and genotype level [bottom right].

**Fig. 4:**
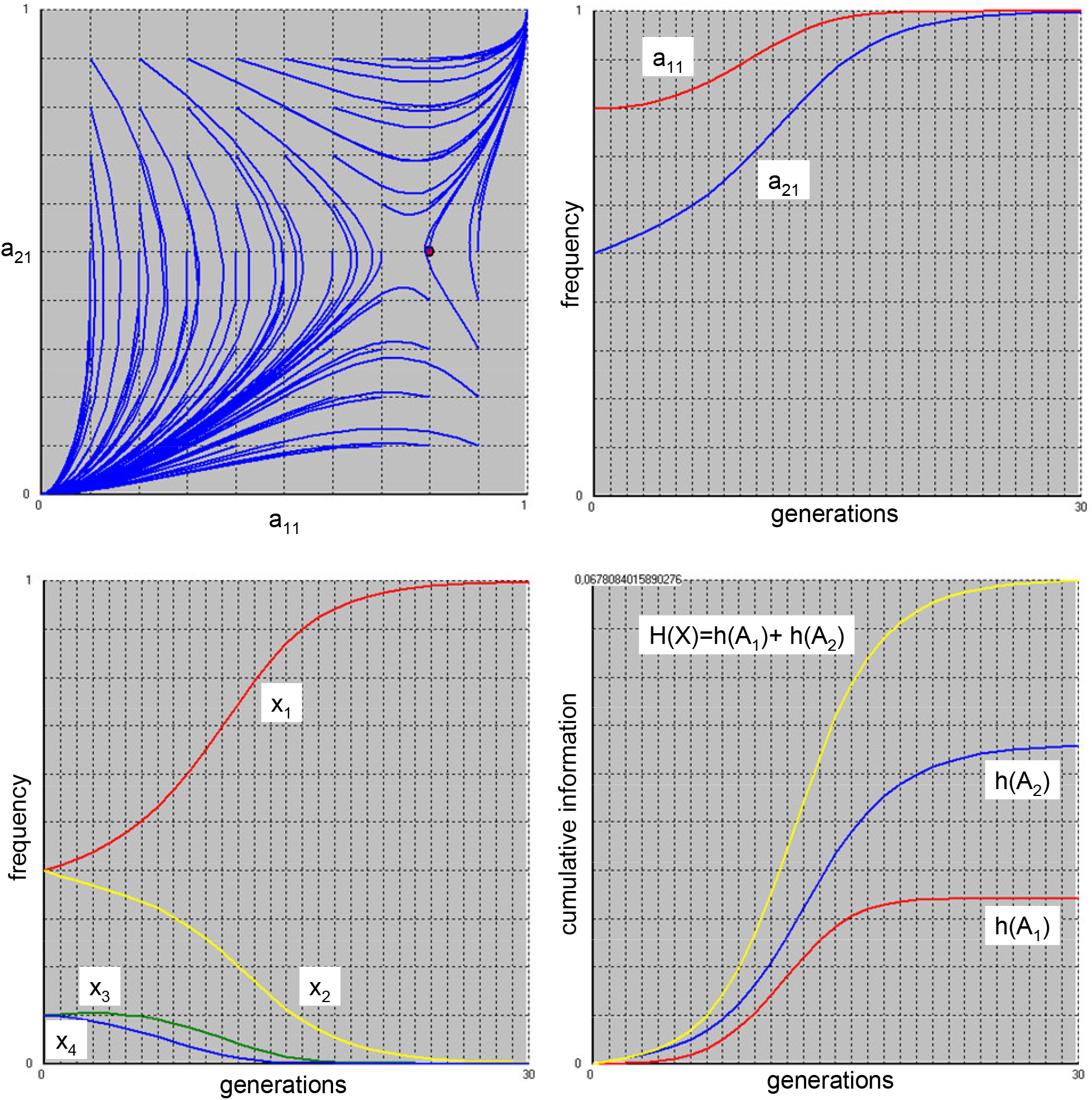
Selection **with recombination** in a two gene loci, two alleles per locus model **without additive fitness**. From left to right and top to bottom: [top left] change of frequency of A_11_ and A_21_ alleles shown against each other from different initial frequencies or [top right] shown in time (generations) from initial a_11_=0.8 and a_21_=0.5. Frequencies of the genotypes changing in time (generations) [bottom left]. Increase of information content due to selection on gene and genotype level [bottom right].

The fitness landscape is given by: w(A_11_)=0.1; w(A_12_)=0.3; w(A_21_)=0.2; w(A_22_)=0.4. We suppose that no recombination occurs (Fig. 1). At the beginning the frequencies a_11_ and a_21_ are high but due to low fitness with time both alleles get lost and the superior ones (A_12_ and A_22_) survive. Consequently, only one genotype, X_4_, survives (w(X_1_:A_11_A_21_)=0.3, w(X_2_:A_11_A_22_)=0,5, w(X_3_:A_12_A_21_)=0.5, w(X_4_:A_12_A_22_)=0.7). In this example, where recombination doesn’t take place, increase of information at the genotype level is not the sum of the ones at gene level. For now, because the theory doesn’t yet bridge the gap to thermodynamics, we need a preliminary unit that we call Selbit (the unit of entropy is Joules/Kelvin) and that hopefully will be unnecessary some day. The increase of the cumulative information is 0,34 Selbit per individual.

What changes due to recombination (Fig. 2)? In a population where recombination occurs, there doesn’t change much compared to the population without if fitness is additive. In our example A_11_ and A_21_ get lost because of lower fitness and so only one genotype, X_4_ survives. But here H(X)=h(A1)+ h(A2). Thus, information behaves additive. However, starting with the identical initial and ending with the same final conditions leads to a similar increase of cumulative information (0,34 Selbit per specimen).

Things are more interesting if the fitness-landscape is not additive, but, e.g. saddle-shaped: w(X_1_)=0.4; w(X_2_)=0.2; w(X_3_)=0.3; w(X_4_)=0.5. If there is no recombination the fittest genotype X_4_ will always survive, at least if initially no allele has a frequency of zero. Depending on the initial conditions, at the gene level the frequency change of the alleles sometimes is seemingly odd (Fig. 3), even loops are possible. The reason is that in the a_11_, a_21_ plane the same point can represent different genotype frequencies. However if we start with a_11_=0.8 and a_21_=0.5 no loop occurs. The cumulative information content at genotype level is not the sum of the ones at gene level and the cumulative information increase is 0,19 Selbit per individual.

Taking the same saddle-shaped fitness-landscape but this time with recombination changes a lot. Now it depends on the initial allele frequencies whether the fittest genotype survives (Fig. 4). If we choose initially a_11_=0.8 and a_21_=0.5 this is not the case. Instead the second best survives and the alleles it consists of. The increase of cumulative information content is lower than in the model without recombination, only 0,07 Selbit. However, ΔH can again be computed as the sum of contributions of each gene locus.

### Information change, natural selection, recombination and mutation

“The fundamental problem of communication is that of reproducing at one point either exactly or approximately a message selected at another point”, Shannon^23^ wrote in the introduction to his information theory. If we replace “point” by “time” or “generation” we can use his model to explain mutation: simplified, a message is sent from an information source to the destination. Because of the influence of a “noise source” the received signal may be different from the sent signal. In our case, the message is the genome that is sent to the offspring. The noise source are copying errors and thermodynamic effects (Schrödinger^24^ recognized as early as 1944 that in the microcosm local strong thermal fluctuations occur that can destroy parts of a molecule and argued that this could be the reason of mutations). In our case the noise is called mutation. From the view of information theory it is an unpleasant change of the original message and hence a loss of information. According to Shannon the information loss due to noise is equal to the amount of information that is necessary to reconstruct the original message. This is an insight we shall use later on.

There are different kinds of mutations; Denver et al.^3,4^ recognized that insertions are as common as base replacements and deletions are common as well. Other kinds of mutations don’t occur so often. Within the scope of the present theory, fortunately not the kind, but just the effect on the fitness of a mutation is of importance.

To get some insight in how pressure from deleterious mutations changes information content we use the simple two gene loci, two alleles model with additive fitness (here the assumption of additive fitness is more plausible because the mutants are not adapted to their genetic environment and so a genotype that wears two mutations may well be less fit than one with only one mutation). We determine that both, the rate of mutation and the fitness loss of the mutant are constant. Only the original genes mutate, not the already mutated ones, of course an unrealistic assumption for the sake of simplicity. Mutation rate is defined as the proportion of the frequency of the original allele that mutates per generation. Fitness loss is also seen relative to the origin: Fitness of the original allele is one; that of the mutant is one minus fitness loss. In the first analysis fitness loss for both gene loci is the same. Initially only the original genotype is present and then the forces of mutation, selection and recombination change the population for some thousand generations until some equilibrium is reached. To measure the decrease of information due to mutation we use Shannon’s insight that the information loss is equal to the amount of information that is necessary to reconstruct the original state. We simply stop mutation—but not selection and recombination—and measure the cumulative information increase until the initial conditions are reached again: that is, until the population consists of original genotypes only.

For analysis we change mutation pressure from 0.05 to 0.25 and fitness loss from 0.05 to 0.45, step 0.05. This matrix is analyzed using regression software. The result is very interesting and is shown in Fig. 5.

**Fig. 5:**
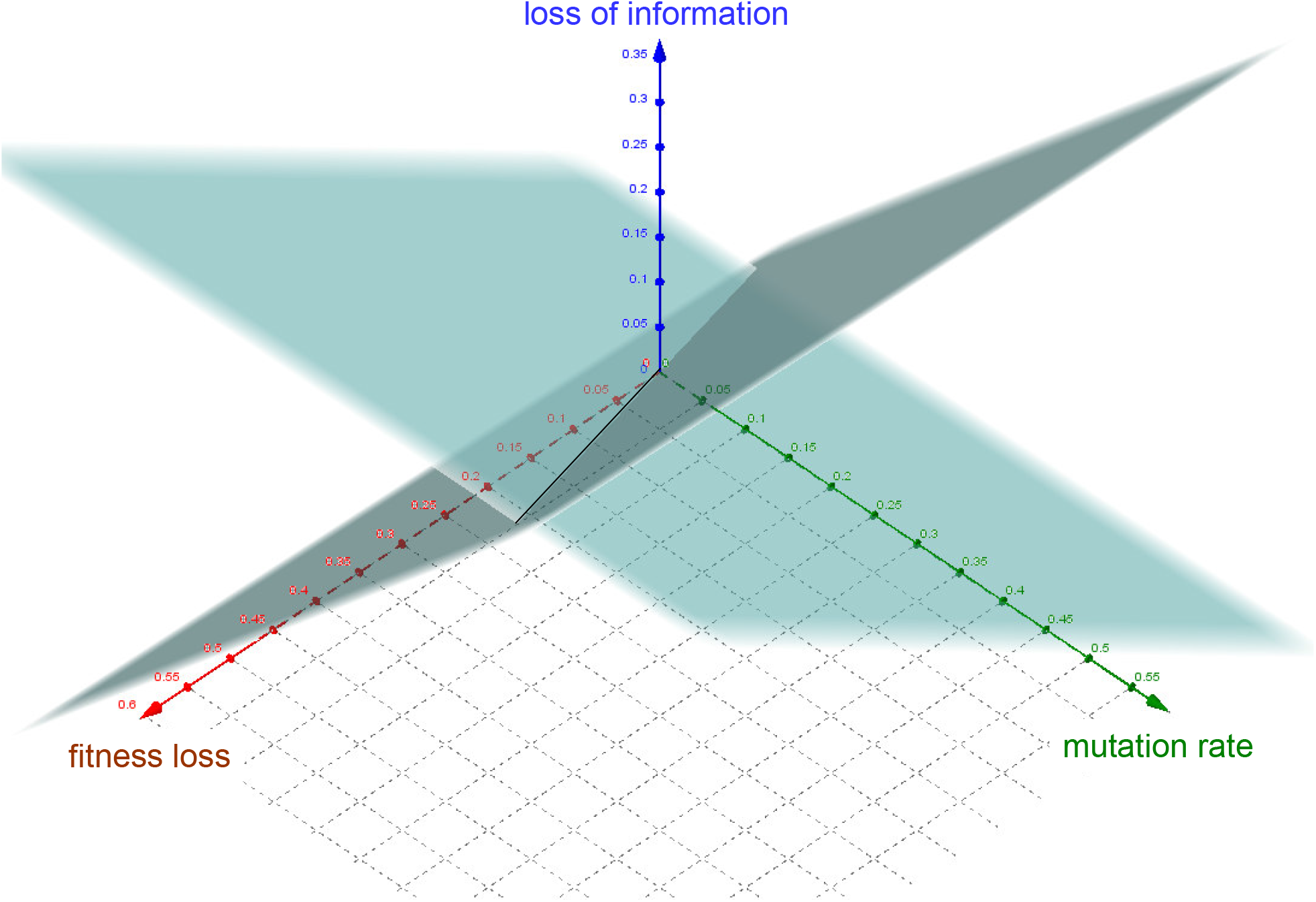
Loss of information due to mutation in a two gene loci, two alleles per locus model with additive fitness. Where there are two values for information loss the lower is the valid one.

Clearly there are two different situations. If selection force is able to compete with mutation rate (dark plane in Fig. 5), information loss depends in a linear way on both, fitness loss and mutation rate and increases with both. If loss of information is z, fitness loss is x and mutation rate is y, equation (40) describes the connection:

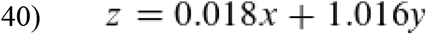

Mutation rate is much more important (about 56 times) than fitness loss. On one hand this means that even if mutants would mutate further and would further loose fitness, the information loss wouldn’t change dramatically as long as the proportion of the original allele in comparison with the collective of mutated ones is in equilibrium. On the other hand this has the consequence that even when fitness loss is very, very small, an information loss proportional to the mutation rate occurs— if not the second situation happens, that we shall discuss in the next paragraph and which is all the more likely, the smaller the fitness loss.

If mutation rate outcompetes selection, so that the original alleles finally disappear, only fitness loss is important to determine information loss:

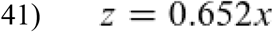

This is the lighter plane in Fig. 5.

Under which circumstances occurs the phase transition? It happens at

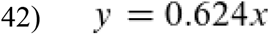

If mutation rate > 0.624*fitness loss, equation (41) describes information loss due to mutation, if mutation rate < 0.624*fitness loss, equation (40).

What happens if the fitness loss of the mutant alleles differs with gene locus? In this case the information loss matches that of a genotype where both mutant alleles have the mean fitness loss even if for one locus y>0.624x and for the other y<0.624x.

With equations (19) and (25) we introduced new tools for the analysis of evolutionary processes that hopefully allow to ask new questions about evolution, e.g.: Is there an upper limit for the acquisition of information due to evolution? Do more gene loci in the genome mean less information increase per locus? What is the meaning of physical conditions, e.g. temperature, for the acquisition of information? Does a change of the environment enhance the acquisition of information? Does cooperation increase information? Do parasitic (selfish) genes decrease information content? And so on. The prerequisite is that empirical data can be used for the calculation of information change. Whole-genome sequencing was used to examine the dynamics of genome sequence evolution of *Saccharomyces cerevisiae*^12^, to study genome evolution of *Escherichia coli* over 50,000 generations^25^ and the long-term adaption to a constant environment^7^. For the last two studies the LTEE (long-term experimental evolution) strains were used^13^. Recently, a barcoding system for the observation of evolutionary dynamics at even higher resolution was established and further developed^14,17^. Thus it is now possible to observe evolution at the gene as well as the genome level and therefore I believe that the view of information theory on evolution could be very helpful and may open new fields of research.

## Supplementary information

*Loss of information due to mutation in a two gene loci, two alleles per locus model with additive fitness. Results of the calculations that led to Fig. 5*.

|  |  | mutation rate/generation |  |  |  |  |
| --- | --- | --- | --- | --- | --- | --- |
| fitness loss |  | 0,05 | 0,1 | 0,15 | 0,2 | 0,25 |
|  | 0,05 | 0,0256 | 0,0256 | 0,0256 | 0,0256 | 0,0256 |
|  | 0,1 | 0,0499 | 0,0526 | 0,0527 | 0,0527 | 0,0527 |
|  | 0,15 | 0,0506 | 0,0812 | 0,0812 | 0,0812 | 0,0812 |
|  | 0,2 | 0,051 | 0,1 | 0,112 | 0,112 | 0,112 |
|  | 0,25 | 0,0515 | 0,1018 | 0,1438 | 0,1438 | 0,1438 |
|  | 0,3 | 0,0521 | 0,1029 | 0,1524 | 0,1782 | 0,1782 |
|  | 0,35 | 0,0526 | 0,104 | 0,1543 | 0,2034 | 0,2151 |
|  | 0,4 | 0,0532 | 0,1052 | 0,1562 | 0,2062 | 0,2548 |
|  | 0,45 | 0,0537 | 0,1064 | 0,1582 | 0,209 | 0,2588 |

